# Rational Design of an Arid Plant–Derived Endophytic Consortium Improves Crop Performance under Controlled Conditions

**DOI:** 10.64898/2026.04.29.721730

**Authors:** Salma Mouhib, Khadija Ait Si Mhand, Imad khatour, Nabil Radouane, Mohamed Hijri

## Abstract

Endophytic bacteria from arid medicinal plants represent a promising source of stress-adapted, plant growth–promoting (PGP) microorganisms. Here, we investigated the cultivable endophytic microbiota of *Peganum harmala* using both nutrient-rich and diluted media to maximize taxonomic recovery. Isolates were dominated by Bacillota and Gammaproteobacteria, with higher diversity in roots than in shoots. Venn analysis revealed a shared core fraction between compartments, forming the basis for consortium assembly. Nine representative strains belonging to *Phyllobacterium, Bacillus, Brevibacillus, Burkholderia, Ralstonia*, and *Amycolatopsis* were selected for functional screening. Pairwise antagonism assays showed high compatibility among Bacillus-related strains, whereas certain taxa exhibited inhibitory interactions, guiding rational consortium design. Functional characterization demonstrated complementary PGP traits, including nitrogen-related activity, phosphate, potassium, and silicate solubilization, siderophore and indole-3-acetic acid production, and ammonia production. No single isolate performed optimally across all traits, supporting a consortium-based strategy. A synthetic bacterial consortium (C2), reconstructed from the core endophytic microbiota using compatibility-guided selection, was evaluated in two crop systems. *In vitro* flax germination assays showed accelerated radicle emergence and improved vigor index, particularly with C2. Under greenhouse conditions, C2 significantly enhanced flax shoot and root biomass, root architecture, leaf area expansion, and photosystem II performance in sterile soil. In faba bean under natural soil, C2 increased leaf number (*p* = 0.02) relative to the control. These results indicate that consortia derived from core endophytes of arid medicinal plants can promote plant growth across diverse crops and soil contexts, although effects remain context-dependent and require rigorous field validation.

**IMPORTANCE:** Endophytic bacteria can serve as sustainable bioinoculants to enhance crop performance under stress conditions. This study demonstrates that the core microbiota of the arid medicinal plant *Peganum harmala* can be rationally assembled into a functionally complementary consortium that improves germination, nutrient acquisition, and whole-plant physiological performance in flax and faba bean. By combining compatibility-guided assembly with functional screening, we show that consortium-based strategies may outperform single-strain inoculants. These findings provide insight into the development of scalable, plant growth–promoting microbial consortia and highlight the importance of testing microbial inoculants under multiple environmental contexts to ensure consistent benefits.

## INTRODUCTION

Growing concerns over soil degradation, excessive fertilizer use, and climate-driven environmental stress are intensifying the need for sustainable strategies to maintain crop productivity. In this context, microbial-based solutions have emerged as promising alternatives to reduce chemical inputs while enhancing plant resilience and nutrient use efficiency (1, 2). Among these, endophytic bacteria, non-pathogenic microorganisms that colonize internal plant tissues, have attracted increasing attention due to their central role in plant health, productivity, and stress tolerance (3).

Unlike rhizospheric or phyllospheric microbes, endophytic bacteria inhabit protected ecological niches within plant tissues, where they can establish stable and persistent associations with their hosts (4–6). This intimate interaction enables them to directly modulate nutrient acquisition, phytohormone balance, and immune signaling pathways, thereby enhancing plant growth and resilience under fluctuating environmental conditions (7–9). Mechanistically, endophytes promote plant performance through multiple pathways, including biological nitrogen fixation, solubilization of inorganic phosphate and potassium, siderophore-mediated iron acquisition, and the production of phytohormones such as indole-3-acetic acid (IAA) (8, 9). In addition, many strains produce antimicrobial compounds and hydrolytic enzymes that suppress phytopathogens and induce systemic resistance, further contributing to plant health (10, 11). Collectively, these traits enhance germination, root development, nutrient uptake, photosynthetic efficiency, and overall plant performance.

Medicinal plants represent particularly valuable reservoirs of functionally diverse endophytic bacteria (9, 12). These plants produce a wide array of bioactive secondary metabolites and often thrive under harsh environmental conditions such as drought, salinity, and nutrient limitation. Such selective pressures favor the recruitment of stress-adapted microbial communities with enhanced metabolic versatility (13–15). Increasing evidence suggests that endophytes associated with medicinal plants not only contribute to plant growth but may also influence the biosynthesis of host secondary metabolites, highlighting their ecological and biotechnological importance (16–18). Consequently, medicinal plants from arid and semi-arid ecosystems constitute underexplored yet promising sources of robust plant growth–promoting (PGP) bacteria (9).

Among these, *Peganum harmala* L., a perennial medicinal plant widely distributed across North African, Middle Eastern, and Central Asian drylands, represents a compelling model system (19). This species is well adapted to alkaline, nutrient-poor soils and water-limited environments, and harbors a diverse endophytic microbiota shaped by long-term exposure to abiotic stress. Previous studies have reported the presence of cultivable endophytes from *P. harmala*, including members of the genera *Bacillus, Pseudomonas*, and *Peribacillus*, many of which exhibit multiple plant growth–promoting traits (20, 21). Similar findings in other arid and halophytic plant species further support the concept that desert medicinal plants are reservoirs of agriculturally relevant microbial diversity (12, 19, 22–24).

Despite the well-documented benefits of individual strains, the application of single-strain inoculants often results in inconsistent performance under field conditions. Environmental variability, competition with native microbiota, and limited ecological fitness frequently constrain the establishment and persistence of introduced strains. As a result, increasing attention has shifted toward the development of multi-strain microbial consortia, or synthetic communities (SynComs), designed to enhance functional robustness and ecological stability (25–27).

Recent advances in microbial ecology highlight that the success of such consortia is fundamentally governed by microbial interactions operating at multiple levels. As emphasized in the recent review by Y. Xu et al. (28), both “within-community” interactions among consortium members and “cross-community” interactions with native soil microbiota critically determine inoculant establishment, stability, and functionality. Positive interactions such as metabolic complementarity and cooperative nutrient exchange can enhance consortium performance, whereas antagonistic interactions may destabilize community structure and reduce efficacy (29). Moreover, native soil communities often exhibit resistance to invasion, limiting the establishment of introduced inoculants. These insights underscore the necessity of integrating ecological theory into the rational design of microbial consortia, including the selection of compatible strains, the promotion of beneficial interactions, and the consideration of ecological fitness in complex soil environments (28).

In this context, compatibility-guided assembly of synthetic consortia represents a promising strategy to improve inoculant performance. By combining metabolically complementary strains while minimizing antagonistic interactions, such approaches aim to enhance functional redundancy, stability, and resilience across diverse environmental conditions. However, key challenges remain, including the identification of compatible taxa, the integration of core microbiota concepts, and the validation of consortium performance across different plant species and soil contexts (6, 8, 9). Moreover, challenges related to formulation, shelf-life stability, and regulatory approval continue to constrain large-scale application (30). Furthermore, translating laboratory findings into agronomic applications requires systematic pipelines encompassing isolation, functional screening, interaction testing, and multi-scale validation (31).

Endophytic bacteria from arid medicinal plants, particularly *P. harmala*, provide a unique opportunity to address these challenges. Their ecological adaptation to stress, combined with multifunctional PGP capacities, makes them ideal candidates for the development of next-generation bioinoculants tailored to marginal environments S. Mouhib et al. (19). Building on these concepts, the present study aimed to systematically isolate and characterize cultivable endophytic bacteria from *P. harmala* using a culture-dependent approach optimized to maximize taxonomic recovery, and to evaluate their potential as bioinoculants through compatibility-guided consortium design.

We hypothesized that (i) multi-strain consortia would outperform single-strain inoculants in promoting plant growth and physiological performance; (ii) consortia composed of taxonomically diverse and functionally complementary strains would generate synergistic effects through coordinated nutrient mobilization, phytohormone modulation, and stress mitigation; and (iii) compatibility-based selection minimizing antagonistic interactions would enhance consortium stability and consistency across different crop systems. By integrating isolation, functional characterization, antagonism assays, and plant performance evaluation, this study provides a framework for bridging laboratory-scale screening with scalable agricultural applications, advancing the development of resilient microbial consortia for climate-smart and sustainable agriculture.

## RESULTS

### Diversity and structure of the cultivable endophytic bacteria

Isolation of endophytic bacteria from *P. harmala* using two different culture media at two different concentrations (TSA, TSA 1/10, PDA, and PDA 1/10) recovered a broad fraction of the cultivable microbiota, although PDA is typically used for fungal isolation. The use of both nutrient-rich and diluted media was associated with increased isolate diversity, underscoring the influence of medium composition on cultivability and diversity recovery. Taxonomic analysis at the phylum and class levels showed that isolates were primarily affiliated with Bacillota and Gammaproteobacteria, with distinct distributions between root and shoot compartments (Figure 1a). The number and composition of taxa differed between roots and shoots (Figure 1b). Venn diagram analysis identified both compartment-specific and shared taxa, including a subset recovered across all culture media (Figure 1c-d). These shared taxa were defined as the cultivable core microbiota of *P. harmala* and were selected for downstream consortium design.

**Figure 1.**
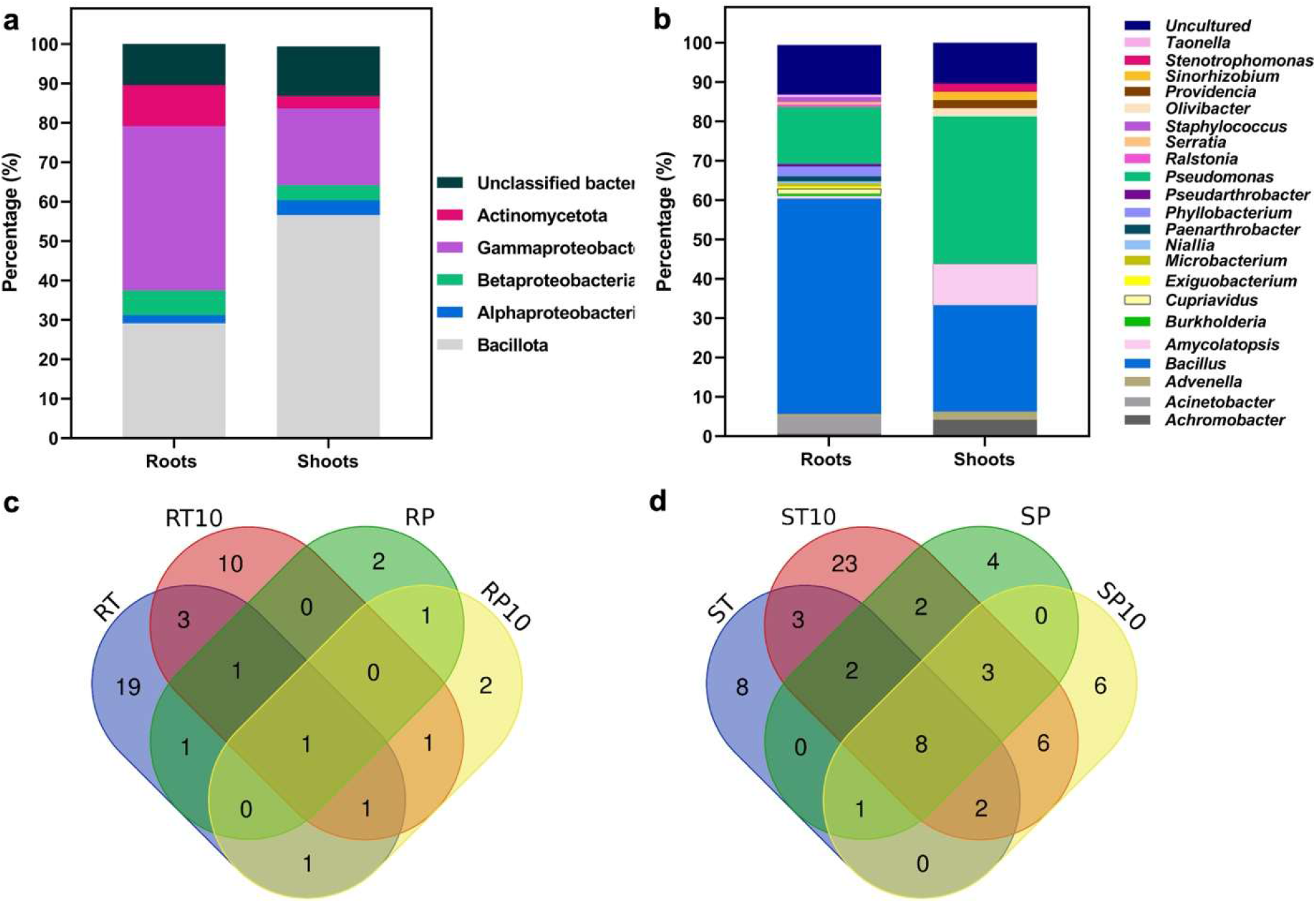
Taxonomic structure, diversity, and distribution of the cultivable endophytic microbiota isolated from roots and shoots of *Peganum harmala*. (a) Distribution at the phylum/class level; (b) distribution at the genus level; (c) number of shared and unique bacterial taxa isolated across different culture media (TSA (T), TSA 1/10 (T10), PDA (P), and PDA 1/10 (P10)) in roots (R); (d) number of shared and unique bacterial taxa in shoots (S).

### Selection and compatibility of core taxa

From the shared root–shoot bacterial community, nine representative isolates were selected for subsequent characterization: *Phyllobacterium myrsinacearum, Bacillus paralicheniformis, Bacillus safensis, Bacillus pumilus, Brevibacillus laterosporus, Burkholderia* sp., *Ralstonia* sp., *Amycolatopsis* sp., and *Bacillus halotolerans*. These strains encompass phylogenetically and functionally diverse taxa, many of which are known for their plant-associated beneficial traits.

Pairwise antagonism assays demonstrated a high degree of mutual compatibility among *Bacillus*-related isolates (*B. paralicheniformis, B. safensis, B. pumilus*, and *B. halotolerans*). In contrast, *Burkholderia* sp., *Ralstonia* sp., and *Brevibacillus laterosporus* displayed inhibitory effects against multiple co-occurring strains (Figure S1). Based on the compatibility profiling, isolates exhibiting minimal antagonistic interactions were preferentially selected for the construction of synthetic consortia, with the aim of maximizing functional complementarity while minimizing inter-strain inhibition (Table 1).

**Table 1.** List of selected endophytic bacteria isolated from both root and shoot tissues of *P. harmala* and compatibility matrix based on pairwise confrontation assays. (+) indicates compatibility between bacterial isolates; (–) indicates inhibitory interaction; NA, not applicable (self-interaction within the same species).

| Bacteria name | *P. myrsinacearum* | *B. paralicheniformis myrsinacearum* | *B. safensis myrsinacearum* | *B. pumilus myrsinacearum* | *B. laterosporus myrsinacearum* | *Burkholderia sp.* | *Ralstonia sp.* | *Amycolatopsis* | *B. halotolerans* |
| --- | --- | --- | --- | --- | --- | --- | --- | --- | --- |
| *Phyllobacterium myrsinacearum* | NA | + | + | + | + | – | + | + | + |
| *Bacillus paralicheniformis* | + | NA | + | + | + | + | + | + | + |
| *Bacillus safensis* | + | + | NA | + | – | – | + | + | + |
| *Bacillus pumilus* | + | + | + | NA | – | – | + | + | + |
| *Brevibacillus laterosporus* | + | – | + | – | NA | – | – | – | – |
| *Burkholderia sp.* | + | – | – | – | – | NA | + | – | – |
| *Ralstonia sp.* | + | – | – | – | – | – | NA | – | – |
| *Amycolatopsis sp.* | + | – | + | + | – | – | – | NA | – |
| *Bacillus halotolerans* | + | + | + | + | + | + | + | + | NA |

### Functional characterization of plant growth–promoting traits

Screening for plant growth–promoting (PGP) traits revealed considerable variability among the selected isolates. Nitrogen-related activity was strongest in isolate *Phyllobacterium myrsinacearum*, whereas potassium solubilization was highest in isolates *B. paralicheniformis* and *Amycolatopsis* sp. (Table 2). Phosphate solubilization was notable in isolate *B. halotolerans*, siderophore production ranged from moderate to strong among *Bacillus* spp., and IAA production was detected in most isolates, with enhanced levels in *B. halotolerans*. Ammonia production was particularly high in the isolate *Amycolatopsis* sp. Importantly, no single isolate displayed universally superior performance across all traits, supporting the rationale for designing multi-strain consortia to exploit functional complementarity (Table 3).

**Table 2.**
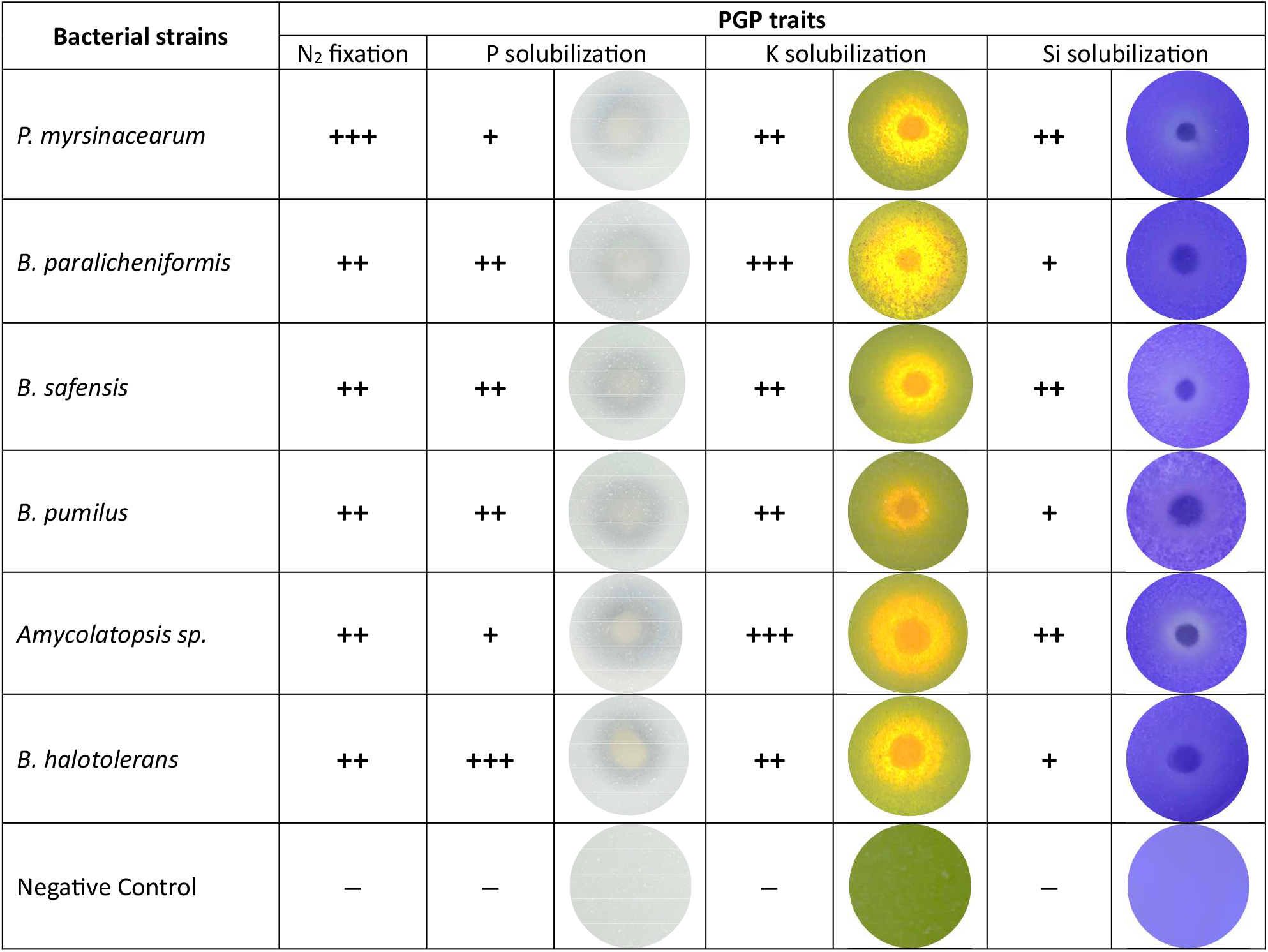
Characterization of plant growth–promoting (PGP) traits among selected core taxa isolates after antagonism assays, including nitrogen (N_2_) fixation and the solubilization of phosphate (P), potassium (K), and silicon (Si). (+++) denotes high expression of the measured trait; (++) denotes intermediate expression; (+) denotes moderate expression; (–) indicates absence or a negative result for the assessed trait.

| Bacterial strains | PGP traits |  |  |  |  |  |
| --- | --- | --- | --- | --- | --- | --- |
|  | N<sub>2</sub> fixation | P solubilization |  | K solubilization |  | Si solubilization |
| *P. myrsinacearum* | +++ | + |  | ++ |  | ++ |
| *B. paralicheniformis* | ++ | ++ |  | +++ |  | + |
| *B. safensis* | ++ | ++ |  | ++ |  | ++ |
| *B. pumilus* | ++ | ++ |  | ++ |  | + |
| *Amycolatopsis sp.* | ++ | + |  | +++ |  | ++ |
| *B. halotolerans* | ++ | +++ |  | ++ |  | + |
| Negative Control | – | – |  | – |  | – |

**Table 3.**
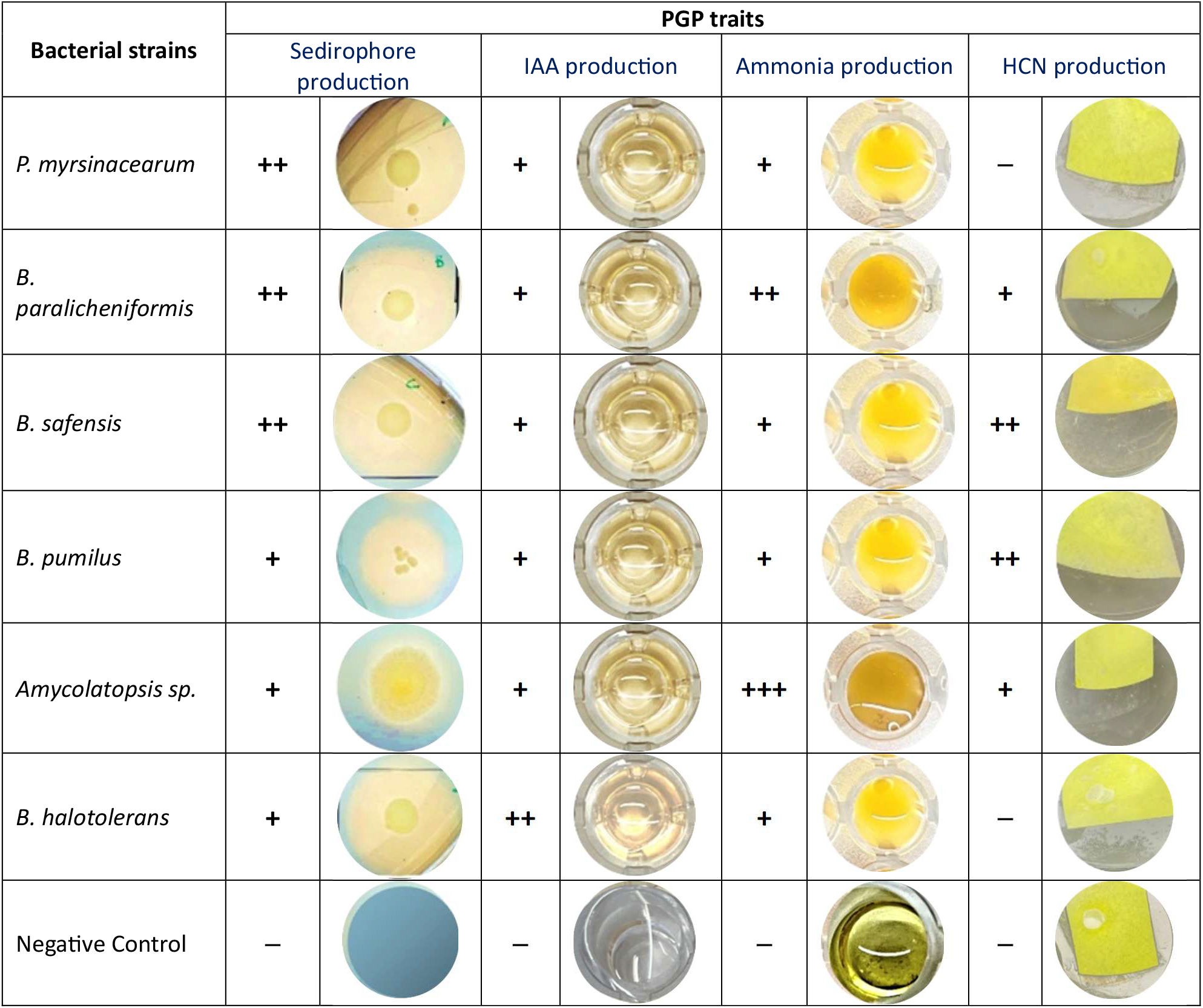
Additional plant growth–promoting (PGP) traits of selected core taxa isolates, including siderophore production, indole-3-acetic acid (IAA) production, ammonia production, and hydrogen cyanide (HCN) production. (+++) denotes high expression of the measured trait; (++) denotes intermediate expression; (+) denotes moderate expression; (–) indicates absence or a negative result for the assessed trait.

### Inoculation effects on flax seed germination and early plant growth

*In vitro* germination assays of flax seeds (*Linum usitatissimum* L. (lin)) on Petri dishes with M medium demonstrated that consortium treatments accelerated germination relative to the controls (Figure 2a). Inoculated seeds exhibited faster radicle emergence and cotyledon expansion within 24–72 h (Table 4). Cumulative germination rates were significantly higher for consortia C2 (*p* = 0.009) and C3 (*p* = 0.01) compared to the positive control (commercial inoculant used for comparison), and vigor index analysis confirmed enhanced early seedling development, with C3 showing the greatest improvement (Figure 2).

**Table 4.**
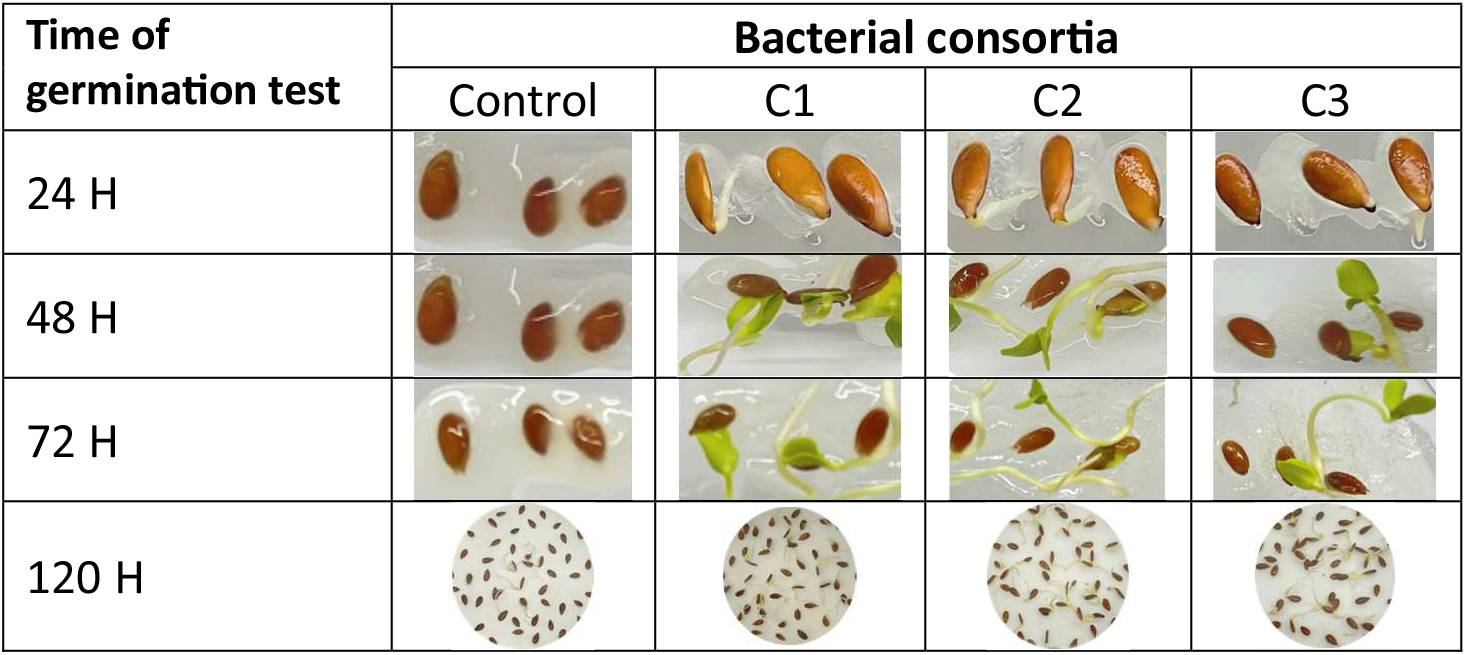
Germination assay of flax seeds conducted in M medium (composed of PhytaGel and mineral nutrients) and in Petri dishes containing distilled water, for the three consortia (C1, C2, and C3) in comparison with the control treatment. Germination was monitored hourly throughout the incubation period.

**Figure 2.**
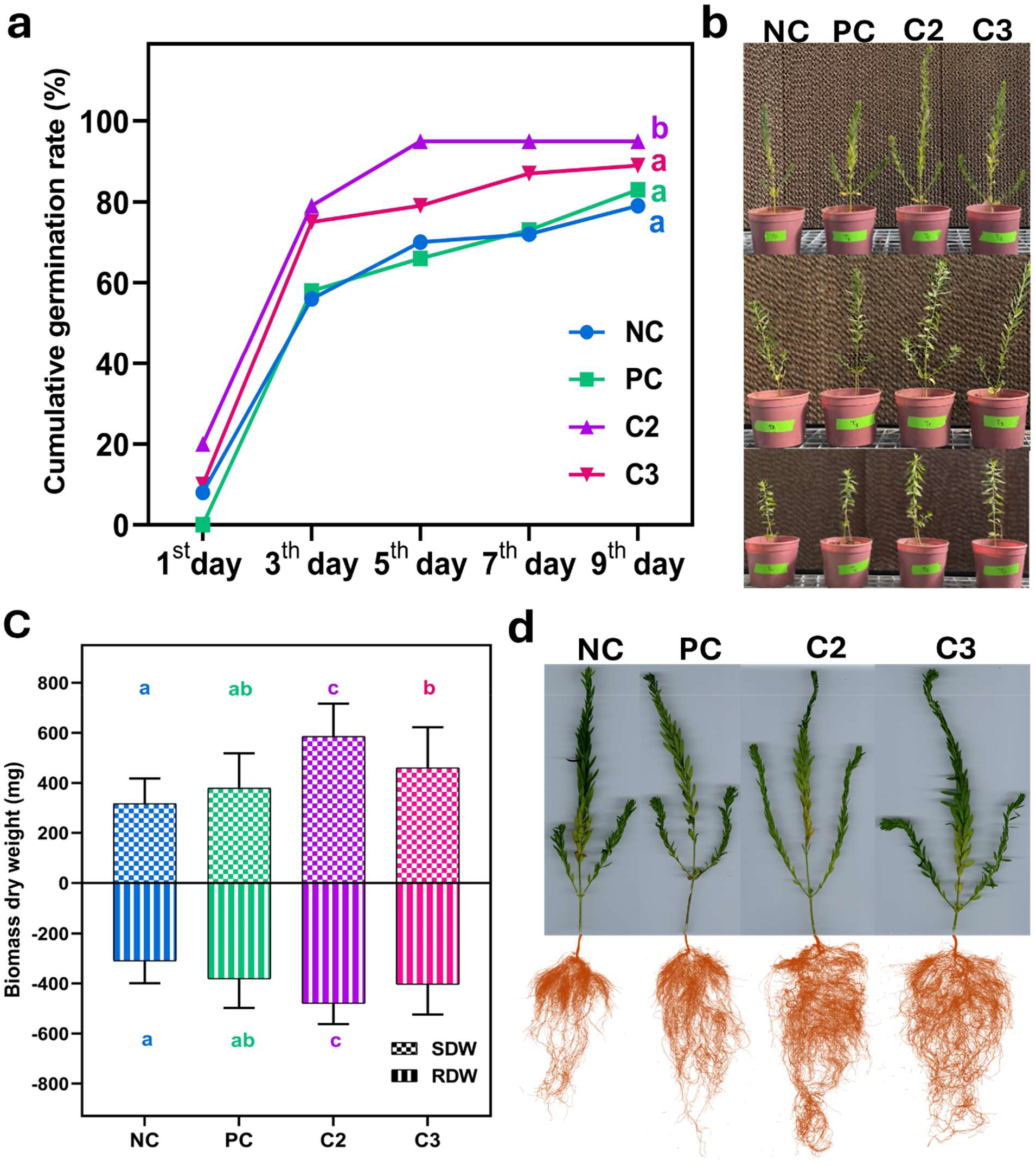
Greenhouse assessment of germination dynamics and growth performance of flax plants inoculated with selected core microbiota consortia. (b) Germination percentage over the first 10 days following treatment application. (c) Representative images of flax plants at 20-day intervals, arranged from bottom to top for each treatment. (c) Bar charts showing shoot and root dry biomass, along with architectural traits of foliar and root systems under greenhouse conditions after consortium inoculation. (d) Representative examples illustrating foliar and root architecture across treatments. NC: negative control; PC: positive control (commercial inoculant); C2 and C3: bacterial consortia developed in this study.

### Effects of inoculation on greenhouse growth performance of flax plants

Under greenhouse conditions, inoculation with the two bacterial consortia significantly enhanced vegetative growth (p = 0.0001), particularly in the C2 treatment (Figure S2). Shoot and root dry weights were highest in C2-treated plants, indicating increased biomass accumulation and enhanced belowground allocation (Figure 2c). Root architecture analysis using WinRHIZO revealed increases in total root length, surface area, and branching (forks) in C2-treated plants, suggesting improved soil exploration and absorptive capacity (Figure 2). Foliar morphometric analysis using WinFOLIA demonstrated increased leaf area and horizontal width, with higher perimeter values reflecting greater leaf expansion rather than shape distortion, indicative of coordinated shoot–root growth enhancement (Figure 3a–c).

**Figure 3.**
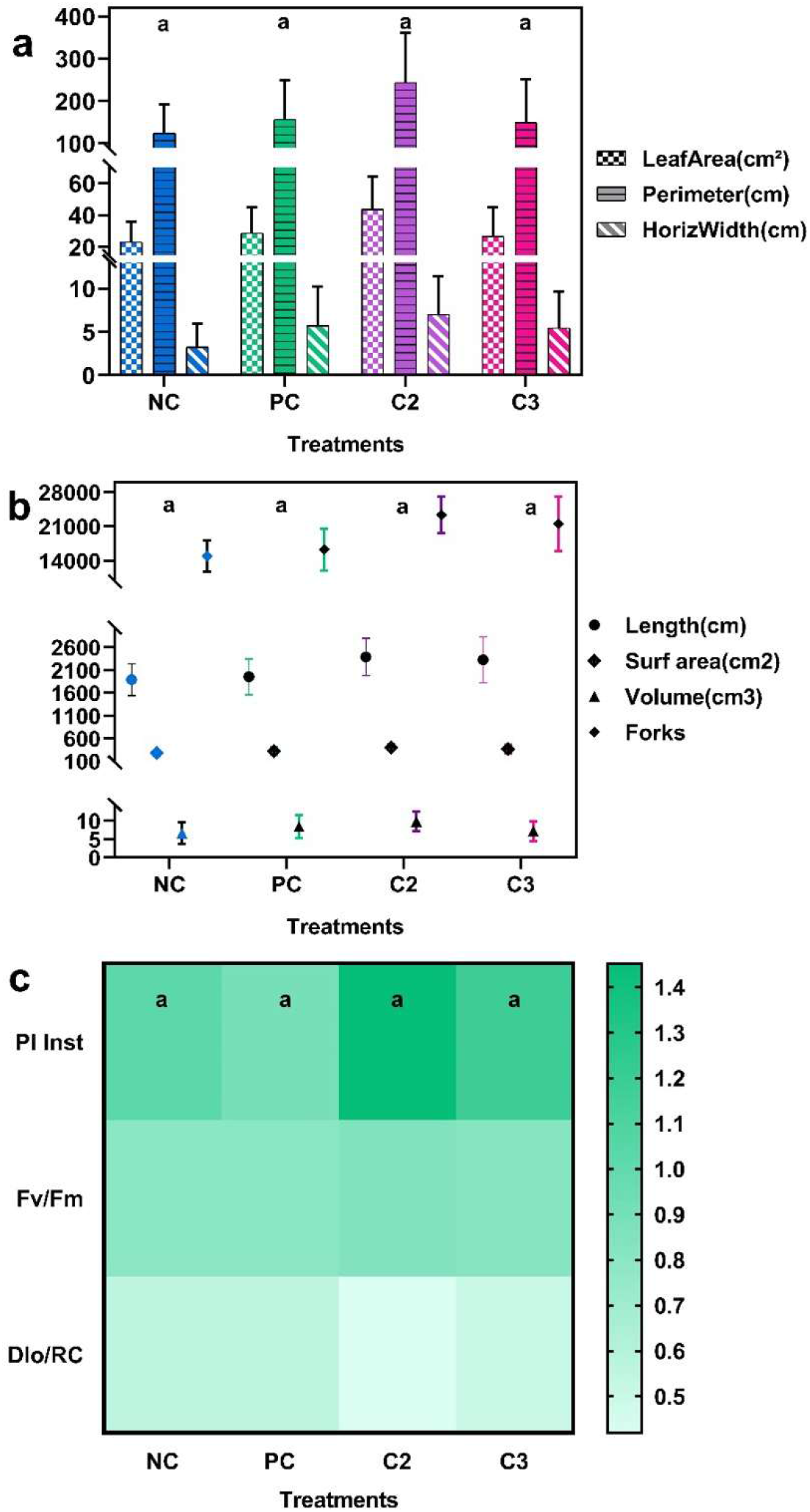
Foliar - root morphometric traits and chlorophyll fluorescence responses of flax plants inoculated with selected consortia. **a.** Foliar parameters (leaf area, perimeter, horizwidth) ascend post-harvest. **b**. Root architecture parameters (length, surface area, volume, and forks) collected post-harvest per treatment. **c**. Chlorophyll fluorescence parameters (PI inst, Fv/Fm, Dio/RC) ascend before-harvest. NC: negative control; PC: positive control (commercial inoculant); C2 and C3: bacterial consortia developed in this study.

### Effects of inoculation on the photosynthetic performance of flax plants

Chlorophyll a fluorescence analysis (using PEA Plus) revealed improved photosystem II (PSII) performance in consortium-treated plants (Figure 3c). The C2 treatment exhibited increased performance index (PI_*abs*_ stable maximum quantum yield of PSII (F_*v*_/F_*m*_),and decreased dissipated energy per reaction center (DI_0_/RC), indicating enhanced photochemical efficiency and reduced energy dissipation. The observed improvement in electron transport efficiency corresponded with increased leaf area and biomass accumulation, suggesting that consortium inoculation promotes coordinated physiological and morphological optimization in flax.

### Greenhouse trial of consortium C2 inoculation on faba bean growth

Germination rates were comparable among treatments, with C2 showing a tendency toward higher cumulative germination than the negative control, which consisted of C2 inoculum autoclaved to kill bacteria (NC) (Figure 4a), although differences were not statistically significant. Plant height measured at 30, 60, and 90 days after sowing followed a consistent trend in which C2-treated plants showing the greatest mean height at each time point, particularly during later developmental stages (Figure 4b).

**Figure 4.**
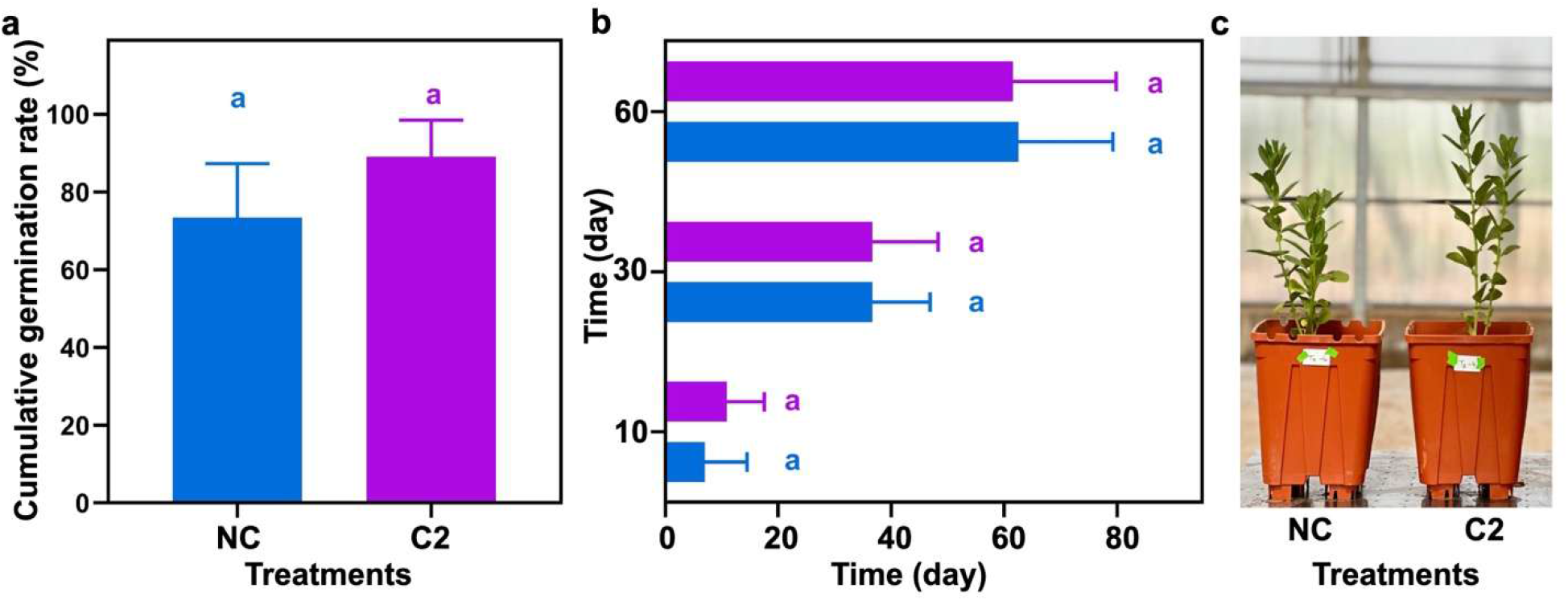
Germination and vegetative growth of faba bean inoculated with consortium C2. (a) Cumulative germination percentage of faba bean seeds following imbibition and soil inoculation with NC and C2 treatments. (b) Plant height (cm) recorded at 30, 60, and 90 days after sowing under the different treatment conditions. (c) Representative photographs of faba bean plants at 60 days (vegetative stage), highlighting phenotypic and morphological differences among treatments.

Morphological assessment revealed that C2-treated plants had higher mean numbers of branches and leaves compared to NC, although these differences were not statistically significant except for leaf number (*p* = 0.020) (Figure S3a). Reproductive parameters, including pod number per plant and pod fresh weight, did not differ significantly between treatments. Cumulative branch and pod production tended to increase in C2-treated plants, although the differences were not significant. Importantly, both C2 and autoclaved C2 (NC) showed comparable baseline performance, confirming the absence of phytotoxicity or antagonistic interactions among the bacterial strains in the C2 consortium (Figure S3a,b).

In addition, spectral indices (NDVI, MCARI, PRI, SIPI) measured at the vegetative stage did not differ significantly among treatments, with values remaining within a narrow range across all experimental groups (*p* > 0.05) (Figure S4a).

## DISCUSSION

Our study demonstrates that endophytic bacteria isolated from *P. harmala* constitute a diverse and functionally versatile microbiota with plant growth–promoting potential and putative contributions to stress resilience. The use of multiple culture media enabled the recovery of a broad spectrum of isolates from both root and shoot compartments, ensuring that the selected core taxa represent a functionally relevant microbial subset. This strategy, based on shared taxa across plant niches, facilitated the identification of strains suitable for synthetic consortium design, combining complementary plant growth–promoting (PGP) traits while limiting potential inter-strain incompatibility.

Importantly, compatibility assays revealed that endophytic bacteria co-occurring within the same host may display antagonistic interactions under *in vitro* conditions. This indicates that spatial co-occurrence does not necessarily reflect functional compatibility, highlighting the importance of systematic antagonism screening prior to consortium assembly (28).

Consortium C2 consistently outperformed other treatments in promoting flax seed germination, root architecture development, leaf expansion, photosynthetic efficiency, and biomass accumulation. The coordinated enhancement of these traits suggests a broad physiological modulation rather than isolated trait-specific effects. This pattern likely reflects complementary mechanisms among strains, including nutrient mobilization, phytohormone production, siderophore and ammonia synthesis, and enzymatic activities that collectively influence root– shoot functioning. These results are consistent with previous reports on endophytes from arid medicinal plants (12, 20–23).

The taxonomic and functional profiles of *P. harmala* endophytes align with earlier studies reporting the predominance of drought-adapted genera such as *Bacillus, Peribacillus*, and *Pseudomonas*, which are frequently associated with antagonistic activity and multiple PGP traits, including phosphate solubilization, siderophore production, IAA synthesis, ammonia and HCN production, and hydrolytic enzyme activity (15, 20, 21). Experimental evidence from related systems further indicates that such isolates can enhance germination, seedling vigor, nutrient acquisition, and biocontrol capacity, particularly under water-limited conditions (12, 23, 32).

Most previous studies on *P. harmala* have focused on single-strain applications or diversity surveys, with limited emphasis on defined multi-strain consortia. In contrast, our results provide evidence that compatibility-filtered consortia can outperform individual strains in promoting plant growth and physiological performance. The inclusion of antagonism screening was critical to ensure functional complementarity, in agreement with consortium design strategies reported for endophytes from other arid ecosystems (25, 26, 33).

While greenhouse experiments confirmed the beneficial effects of C2 (and, to a lesser extent, C3), their performance is expected to be context-dependent under field conditions. Soil physicochemical properties, plant genotype, environmental stress, and agronomic practices are known to influence inoculant efficacy, underscoring the need for multi-environment validation (12, 25, 34). In addition, biosafety evaluation remains a necessary step prior to field deployment.

The agronomic performance of consortium C2 was further evaluated in a distinct crop system (faba bean, *Vicia faba* L.). Although physiological spectral indices showed no significant differences among treatments, C2 consistently tended to increase leaf number, germination rate, plant height, and branching. These trends suggest a moderate growth-promoting effect. The comparable performance of the autoclaved control further indicates that, under non-stressful and relatively favorable conditions, plant performance may be primarily governed by substrate fertility and environmental stability rather than microbial activity alone.

Plant–microbe interactions are often more pronounced under nutrient limitation or abiotic stress (35, 36). Accordingly, the absence of strong statistical separation does not negate the biological potential of consortium C2 but rather highlights the strong context dependency of microbial effects..

Additionally, the limited response in faba bean may be partly explained by its leguminous nature and symbiosis with rhizobia, which enables nitrogen fixation through nodule formation. This endogenous nitrogen acquisition may reduce the relative contribution or detectability of additional plant growth–promoting bacteria under the tested conditions.

A key outcome of this study is the evaluation of consortium performance across contrasting experimental contexts. In flax, plants were grown under controlled sterile conditions, whereas the faba bean experiment was conducted in non-sterile soil under more complex microbial background conditions. Despite these differences, C2 maintained consistent growth-promoting trends, suggesting functional robustness across environments. This stability is consistent with recent reports highlighting the ecological resilience of plant-derived microbial consortia (37–39).

Interestingly, the response observed with the autoclaved consortium suggests that the formulation matrix may also contribute to plant performance, potentially through residual metabolites or soil-conditioning effects. This could enhance formulation stability and broaden potential applications of microbiome-based biostimulants.

From an ecological perspective, the cross-species activity of C2 supports the hypothesis that core endophytic taxa possess broadly conserved plant growth–promoting functions beyond their original host. The consistency observed between flax and faba bean further supports a holobiont-informed, reductionist consortium design approach (40).

Collectively, these findings indicate that compatibility-guided synthetic consortia derived from core microbiota represent a promising strategy for biostimulant development. Such consortia can achieve performance comparable to commercial inoculants while ensuring functional stability and absence of phytotoxic effects. Their validation across distinct plant systems further highlights their translational potential for sustainable crop management.

Overall, our results provide partial support for the proposed hypotheses. First, multi-strain consortia, particularly C2, generally outperformed individual strains in promoting plant growth and physiological traits, supporting hypothesis (i). Second, the coordinated improvements observed across multiple plant parameters are consistent with the expected synergistic effects of taxonomically diverse and functionally complementary strains, lending support to hypothesis (ii), although direct mechanistic validation was not assessed. Finally, the stable performance of C2 across contrasting experimental systems, combined with the importance of antagonism screening during consortium assembly, supports hypothesis (iii), suggesting that compatibility-based selection contributes to functional stability. However, the context-dependent responses observed, particularly under non-stress conditions and in legume systems, indicate that these effects may vary depending on environmental and host-related factors.

## CONSLUSIONS

Our results indicate that compatibility-guided consortia derived from core taxa can enhance plant performance across several traits, including germination, plant architecture, photosynthetic efficiency, and biomass accumulation, while maintaining functional consistency across different plant hosts and growth conditions. Although the magnitude of these effects varied depending on the experimental context, the overall trends support the value of multi-strain, functionally complementary consortia over single-strain approaches. The importance of compatibility-based selection was also evident, contributing to stable and reproducible consortium performance.

These findings highlight the potential of *P. harmala* endophytes as a resource for the development of microbial biostimulants adapted to arid and resource-limited environments. Future work should focus on optimizing consortium design, conducting field validation across diverse crops and environmental conditions, and addressing biosafety and regulatory considerations. Expanding the exploration of endophytes from other arid-adapted plants may further broaden the pool of beneficial microbial candidates.

## MATERIALS AND METHODS

### Sample collection and plant material

Healthy individuals of *Peganum harmala* L. were collected from arid environments during the active vegetative stage (19). Entire plants were carefully uprooted to preserve the root system, placed in sterile bags, and transported to the laboratory under cooled conditions for immediate processing. In the laboratory, roots and shoots were separated aseptically and processed independently for endophyte isolation.

### Isolation of the cultivable endophytic microbiota

Endophytic bacteria were isolated from surface-sterilized roots and shoots following the protocol previously adopted by S. Mouhib et al. (19) with minor modifications. Briefly, plant tissues were thoroughly washed under running tap water to remove adhering soil particles and subjected to sequential surface sterilization using ethanol and sodium hypochlorite, followed by repeated rinsing in sterile distilled water. The efficacy of sterilization was verified by imprinting treated tissues onto agar plates to confirm the absence of epiphytic growth. Sterilized tissues were macerated in sterile saline solution, serially diluted, and plated onto four culture media: Tryptic Soy Agar (TSA), TSA diluted tenfold (TSA 1/10), Potato Dextrose Agar (PDA), and PDA diluted tenfold (PDA 1/10). The combination of nutrient-rich and diluted media was used to capture both fast-growing copiotrophic bacteria and slow-growing oligotrophic endophytes, thereby increasing taxonomic richness and minimizing medium-driven selection bias. Plates were incubated at 28 °C for 48–72 h, and morphologically distinct colonies were purified through repeated streaking and preserved in glycerol stocks at −80 °C.

### Molecular identification and core taxa selection

Genomic DNA was extracted from purified isolates using standard bacterial DNA extraction procedures described in S. Mouhib et al. (19). The 16S rRNA gene was amplified using universal primers pA and 926R, and PCR products were purified and subjected to Sanger sequencing (19). Raw sequences were edited and assembled before comparison against the NCBI GenBank database using both BLASTn and Geneious Prime v2023.0.3 (Biomatters, Auckland, New Zealand) for taxonomic identification. Presence–absence data were compiled across plant compartments and culture media to determine shared taxa. Only isolates recovered from both roots and shoots were considered members of the cultivable core microbiota and were retained for subsequent functional characterization and consortium design.

### Antagonism and compatibility assays

Pairwise compatibility among selected isolates was assessed using confrontation assays on agar plates. Each strain was cross-streaked against all other strains and incubated at 28 °C for 48 h. The presence of inhibition zones was recorded as evidence of antagonistic interactions. Isolates exhibiting mutual tolerance without growth suppression were considered compatible. A compatibility matrix was constructed to guide rational assembly of multi-strain consortia while minimizing inhibitory interactions.

### Characterization of plant growth–promoting traits

Selected isolates were screened for multiple plant growth–promoting (PGP) traits using established qualitative and semi-quantitative assays. Each isolate was cultured in TSB for 24 h at 28° C under agitation (150 rpm), harvested by centrifugation (8000 rpm, 5 min), washed with PBS, and adjusted to OD600 = 0.8. Nitrogen activity was evaluated by streaking isolates onto nitrogen-deficient combined carbon (CC) medium (41), phosphate solubilization was assessed using the National Botanical Institute’s phosphate (NBRIP) growth medium (42), potassium solubilization was tested on Aleksandrov medium containing Feldspath as the potassium source (43), silicon solubilization was evaluated on nutrient agar supplemented with silicate using bromocresol purple as pH indicator (44), siderophore production was assessed onto (Chrome Azurol S agar following a modified protocol of (45), Indole-3-acetic acid (IAA) production by the bacterial isolates was assessed using a qualitative assay, following the method by C. L. Patten and B. R. Glick (46), ammonia production, and hydrogen cyanide (HCN) production were evaluated according the protocols (47, 48). Trait intensity was scored based on halo formation, solubilization index, or colorimetric changes relative to uninoculated controls.

### Consortium design

Consortium formulation was based on three main criteria: inclusion within the shared root–shoot core microbiota, functional complementarity of PGP traits, and compatibility according to the antagonism matrix. Three candidatesynthetic consortia were rationally assembled from mutually compatible core taxa. Consortium C1 comprised *Phyllobacterium myrsinacearum, Bacillus safensis, Bacillus pumilus*, and *Amycolatopsis* sp. Consortium C2 included *P. myrsinacearum, B. paralicheniformis, B. safensis, B. pumilus*, and *B. halotolerans*. Consortium C3 combined *P. myrsinacearum, B. paralicheniformis, B. safensis, B. pumilus, Amycolatopsis* sp. and *B. halotolerans*. Individual strains were cultured in nutrient broth to the late exponential phase, harvested by centrifugation, washed, and adjusted to approximately 10^8^ CFU mL^−1^. Equal volumes of compatible strains were combined to generate each consortium inoculum.

### Germination assays

Seeds of *Linum usitatissimum* L. (lin) variety Zriat l’kettan, were surface-sterilized and immersed in bacterial suspensions (10^8^ CFU mL^−1^) for 2 h, while control seeds were treated with sterile saline solution. For preliminary screening, treated seeds were placed on moistened sterile filter paper in Petri dishes and incubated at 25 °C under a 16 h light/8 h dark photoperiod. Germination percentage and kinetics were recorded daily. Seeds were additionally evaluated on M medium to confirm early germination performance under semi-sterile conditions. Based on germination rate and vigor index, the two best-performing consortia were selected for greenhouse validation.

### Greenhouse experiment with flax plants

A completely randomized design was implemented under greenhouse conditions, with 20 replicates per treatment, in which each pot (5 L) of natural sterilized soil was inoculated with 50 mL of each treatment. Treatments included a non-inoculated control (negative control), selected consortium C2, selected consortium C3, and a commercial rhizospheric biostimulant reference (B1). Plants were grown under uniform irrigation and standard agronomic practices until the pre-flowering stage.

### Measurement of plant growth, physiological, and morphometric parameters

Plant growth and physiological performance were evaluated at the pre-flowering stage using an integrated set of morphological and functional measurements. Chlorophyll *a* fluorescence was recorded on dark-adapted leaves prior to harvest using a Handy PEA fluorimeter (Hansatech Instruments, UK). Fluorescence transients were analyzed with PEA Plus software according to the OJIP test framework (49), and parameters including F_*v*_/F_*m*_ (maximum quantum efficiency of PSII), PIabs (performance index on absorption basis), and DI_0_/RC (energy dissipation per reaction center) were extracted to assess photosynthetic efficiency.

Root systems were carefully washed free of substrate, scanned, and analyzed using WinRHIZO™ software (Regent Instruments Inc., Canada). Root architectural traits quantified included total root length, root surface area, root volume, and the number of forks as an indicator of branching intensity and soil exploration capacity. Aboveground morphometric traits were assessed using WinFOLIA™ software (Regent Instruments Inc., Canada), measuring leaf area, leaf perimeter, and horizontal width to characterize shoot development and canopy expansion. Finally, shoot and root biomass were determined after oven-drying plant material at 65 °C until constant weight, allowing quantification of dry matter accumulation and biomass allocation patterns.

### Greenhouse experiment with faba bean plants

To assess the agronomic performance of the selected bacterial consortium (C2) beyond flax, a greenhouse experiment was conducted using faba bean (*Vicia faba* L.) variety Violette (AGRIN, Morocco) under a randomized complete block design, using natural soil collected from the experimental farm of UM6P, Benguerir (32°13′12.84″N, 7°53′11.78″W). Two treatments were evaluated: NC (negative control), consisting of an autoclaved C2 inoculum (non-viable), and C2, corresponding to the active bacterial consortium.

The bacterial suspension was prepared as described for the flax experiment. Seeds were imbibed for 1 h in the corresponding suspension prior to sowing, and each pot (20 L) containing natural soil received 100 mL of the respective inoculum. Two seeds were sown per pot, and after emergence, one uniform seedling was retained.

The experiment included four blocks with eight replicates per treatment per block. Germination rate was recorded during early establishment. Plant height was measured at 30, 60, and 90 days after sowing. Branch and leaf numbers were recorded during the vegetative stage, and flowering time was monitored to assess phenological responses. At maturity, pod number and pod fresh weight were measured. Physiological indices (NDVI, MCARI, PRI, SIPI) were assessed on fresh leaves using a PolyPen device. Plants were irrigated every two days at approximately half of the substrate water-holding capacity.

### Statistical analysis

Data were tested for normality prior to statistical evaluation. Treatment effects were assessed using one-way analysis of variance (ANOVA) and two-way ANOVA including block as a factor, and mean comparisons were performed using Tukey’s HSD test at p < 0.05. Heatmap visualization and multivariate analyses were conducted to provide an integrative assessment of physiological and growth responses across treatments, and the results were visualized in GraphPad Prism 8.0.1 (244). All statistical analyses were conducted in R (2025.09.2+(418).

N.B: Root nodulation and final dry biomass measurements were not included due to severe environmental stress caused by a heat wave and aphid infestation during the late experimental phase.

## ACKNOWLEDGMENTS

This work was supported by the OCP Nutricrops Project AS-85, whose support is gratefully acknowledged. We also thank Nutribiotek for providing access to their SmartFarm infrastructure for conducting the greenhouse assays. We are grateful to Narjisse Mahmoudi, Abdeljalil Zalarh and Daouda Ndiaye for their valuable assistance and technical support.

## DATA AVAILABILITY

Data Availability Statement: The 16S rRNA gene sequences generated in this study are publicly available in Zenodo under DOI: 10.5281/zenodo.19432075 (https://zenodo.org/records/19432075). Sequences corresponding to the synthetic consortium strains have been deposited in GenBank under accession numbers PZ058640–PZ058648.

